# Interferon lambda signals in maternal tissues to exert protective and pathologic effects in a gestational-stage dependent manner

**DOI:** 10.1101/2022.01.04.475019

**Authors:** Rebecca L. Casazza, Drake T. Philip, Helen M. Lazear

## Abstract

Interferon lambda (IFN-λ, type III IFN) is constitutively secreted from human placental cells in culture and reduces Zika virus (ZIKV) transplacental transmission in mice. However, the roles of IFN-λ during healthy pregnancy and in restricting congenital infection remain unclear. Here we used mice lacking the IFN-λ receptor (*Ifnlr1*^-/-^) to generate pregnancies lacking either maternal or fetal IFN-λ responsiveness and found that the antiviral effect of IFN-λ resulted from signaling exclusively in maternal tissues. This protective effect depended on gestational stage, as infection earlier in pregnancy (E7 rather than E9) resulted in enhanced transplacental transmission of ZIKV. In *Ifnar1*^-/-^ dams, which sustain robust ZIKV infection, maternal IFN-λ signaling caused fetal resorption and intrauterine growth restriction. Pregnancy pathology elicited by poly(I:C) treatment also was mediated by maternal IFN-λ signaling, specifically in maternal leukocytes, and also occurred in a gestational stage-dependent manner. These findings identify an unexpected effect of IFN-λ signaling specifically in maternal (rather than placental or fetal) tissues, which is distinct from the pathogenic effects of IFN-αβ (type I IFN) during pregnancy. These results highlight the complexity of immune signaling at the maternal-fetal interface, where disparate outcomes can result from signaling at different gestational stages.

**IMPORTANCE:** Pregnancy is an immunologically complex situation, which must balance protecting the fetus from maternal pathogens with preventing maternal immune rejection of non-self fetal and placental tissue. Cytokines, such as interferon lambda (IFN-λ), contribute to antiviral immunity at the maternal-fetal interface. We found in a mouse model of congenital Zika virus infection that IFN-λ can have either a protective antiviral effect or cause immune-mediated pathology, depending on the stage of gestation when IFN-λ signaling occurs. Remarkably, both the protective and pathologic effects of IFN-λ occurred through signaling exclusively in maternal immune cells, rather than in fetal or placental tissues, or in other maternal cell types, identifying a new role for IFN-λ at the maternal-fetal interface.

## INTRODUCTION

Immune regulation at the maternal-fetal interface is complex due to conflicting immunological objectives: protection of the fetus from maternal pathogens, and prevention of immune-mediated rejection of the semi-allogeneic fetus and placenta. The few pathogens able to surmount the placental barrier and cause congenital infections include Zika virus (ZIKV), rubella virus (RUBV), and human cytomegalovirus (1). The mechanisms by which pathogens are excluded from the fetal compartment are not fully understood, and it is unclear how antiviral activity at the maternal-fetal interface affects tolerogenic immunity. Moreover, pregnancy encompasses multiple developmental stages including implantation, fetal growth, and parturition, each with unique immunologic requirements (2–4). Because the physiology and immunology of the placenta change over gestation, there likely are distinct antiviral mechanisms at each stage of pregnancy. The need to balance protective and pathogenic immunity is not unique to the maternal-fetal interface: epithelial surfaces such as the gastrointestinal and respiratory tracts encounter microbes and must provide protection from pathogens without inflicting inflammatory damage. Interferon lambda (IFN-λ, type III IFN) is a cytokine that elicits a similar antiviral transcriptional response as type I IFNs (IFN-αβ), but signals through a distinct heterodimeric receptor comprised of IFNLR1 and IL10Rb. (5). The IFN-λ receptor is predominantly expressed on epithelial cells and consequently confers antiviral protection at barrier surfaces including the gastrointestinal and respiratory tracts. IFN-λ is secreted constitutively from human mid-gestation and term placental explants and trophoblasts cultured ex vivo, human trophoblast organoids, and in human placental cell lines syncytialized in culture (6–8). In a mouse model of congenital ZIKV infection, IFN-λ restricted transplacental transmission, as fetuses from *Ifnlr1^-/-^* pregnancies (*Ifnlr1^-/-^* x *Ifnlr1^-/-^*) sustained higher fetal and placental viral loads than those from wild-type pregnancies (9). However, the mechanism by which IFN-λ protects against viral infection at the maternal-fetal interface has not been defined.

The 2015-2016 ZIKV outbreak throughout Latin America and the Caribbean revealed that ZIKV infection during pregnancy can produce a spectrum of adverse fetal and neonatal outcomes (collectively referred to as congenital Zika syndrome) including microcephaly, intrauterine growth restriction (IUGR), placental insufficiency, vision and hearing loss, as well as miscarriage and stillbirth (10, 11). Infants born without overt congenital Zika syndrome also can have cognitive or functional deficits that become evident later in infancy or childhood (12–14). Mouse models of ZIKV congenital infection have been developed to test vaccines and antivirals as well as to define ZIKV pathogenic mechanisms and antiviral immunity at the maternal-fetal interface (15–17). Aspects of ZIKV fetal pathogenesis are recapitulated in mouse models and include fetal loss, IUGR, fetal brain infection, placental pathology, and neurologic defects. The outcomes of congenital ZIKV infection usually are more severe when infection occurs earlier in gestation in both mice (9, 15, 18) and humans (Brady et al., 2019; Hoen et al., 2018; Honein et al., 2017; Ospina et al., 2020). Although there are differences between mouse and human pregnancy (23), mice provide a genetically tractable system to study antiviral and placental immunity at distinct gestational timepoints.

Here we used mouse models of congenital ZIKV infection to determine the targets of IFN- λ signaling by infecting pregnancies that lacked IFN-λ signaling (*Ifnlr1^-/-^*) in maternal and/or fetal tissues. When we infected at embryonic day 9 (E9), we observed that IFN-λ signaling in maternal tissues protected against transplacental ZIKV transmission. Surprisingly, IFN-λ had a deleterious effect when pregnancies were infected two days earlier at E7, with IFN-λ responsive dams exhibiting higher rates of ZIKV transmission as well as overt pathology and fetal resorption. This effect was not specific to ZIKV as we also found that maternal IFN-λ signaling increased rates of fetal loss after poly(I:C) treatment and that this pathology similarly was dependent on gestational age at the time of administration. These findings identify an unexpected effect of IFN-λ signaling specifically in maternal (rather than placental or fetal) tissues and highlight the complexity of immune signaling at the maternal-fetal interface, where disparate outcomes can result from signaling at different gestational stages.

## RESULTS

### ZIKV congenital infection is exacerbated earlier in pregnancy and in Ifnar1^-/-^ dams

ZIKV replication in mice is restricted by the IFN response because ZIKV is unable to antagonize mouse STAT2 (24, 25). Thus, mouse models of ZIKV pathogenesis typically employ mice lacking IFN-αβ signaling, usually through genetic loss of the IFN-αβ receptor (*Ifnar1^-/-^)* alone or in combination with the IFN-γ receptor (*Ifnar1^-/-^ Ifngr1^-/-^* DKO), or by treatment of wild-type mice with an IFNAR1-blocking monoclonal antibody (MAR1-5A3) (26). Congenital ZIKV pathogenesis has been studied in many different mouse models that vary in mouse genetic background, IFN responsiveness, ZIKV strain, inoculation route, duration of infection, and gestational stage at infection and harvest (27). To better define the conditions that produce transplacental transmission and pathology, we evaluated gross pathology and fetal viral loads in pregnant *Ifnar1^-/-^* dams or wild-type dams treated with 2mg of MAR1-5A3 1 day prior to infection. To exclude fetal pathology resulting from severe maternal morbidity, we first compared the virulence of three Asian-lineage ZIKV strains in non-pregnant female 8-10 week-old *Ifnar1*^-/-^ mice infected with 1000 FFU of ZIKV by subcutaneous inoculation in the footpad (Figure 1A and B). We found that strain H/PF/2013 was the most virulent, causing 80% lethality, whereas strain FSS13025 caused modest weight loss in some mice and only 20% lethality, and strain PRVABC59 caused no weight loss or lethality, altogether consistent with prior studies reporting the relative virulence of these strains in *Ifnar1*^-/-^ mice of various ages and inoculation routes (Carbaugh et al., 2020; Lazear et al., 2016; Tripathi et al., 2017). We chose to use strain FSS13025 for further experiments to achieve robust maternal infection without severe maternal morbidity and because of its use in studies from other groups evaluating the role of IFN signaling in congenital ZIKV infection (29). We infected pregnant dams with 1000 FFU of ZIKV FSS13025 by subcutaneous inoculation in the footpad at E7 or E9 (Figure 1C) and measured viral loads in the maternal spleen, placentas, and fetal heads at E15 (8 or 6 days post-infection (dpi)), (Figure 1D-F). We observed higher viral loads in *Ifnar1*^-/-^ dams compared to WT dams treated with MAR1-5A3, and viral loads were higher after infection at E9 (6 dpi) compared to infection at E7 (8 dpi). Placental and fetal viral loads corresponded to maternal spleen viral loads, suggesting that fetal infection increases with the severity of maternal infection. Rates of transplacental transmission (measured by proportion of fetal heads that were ZIKV-positive) were higher in *Ifnar1^-/-^* dams compared to MAR1-5A3-treated dams (64% vs. 57% at E7, 100% vs. 61% at E9). All fetuses that were intact (not resorbed) were photographed and weighed (Figure 1G- I). Fetuses smaller than one standard deviation below the mean of uninfected pregnancies were classified as having IUGR. *Ifnar1*^-/-^ dams exhibited significantly higher resorption rates compared to uninfected controls. In contrast to fetal viral loads, which were higher in dams infected at E9, fetal pathology was greater in dams infected at E7, suggesting higher placental/fetal susceptibility early in pregnancy or that pathology increases with longer infection times. The results were the same when we assessed pathology by crown-rump length (CRL) rather than fetal weight (Figure S1A-B). These results indicate that there are significant differences in adverse pregnancy outcomes when infections occur at different gestational stages, and that fetal pathologic outcomes and viral loads are more severe in the context of high maternal infection (*Ifnar1*^-/-^).

**Figure 1.**
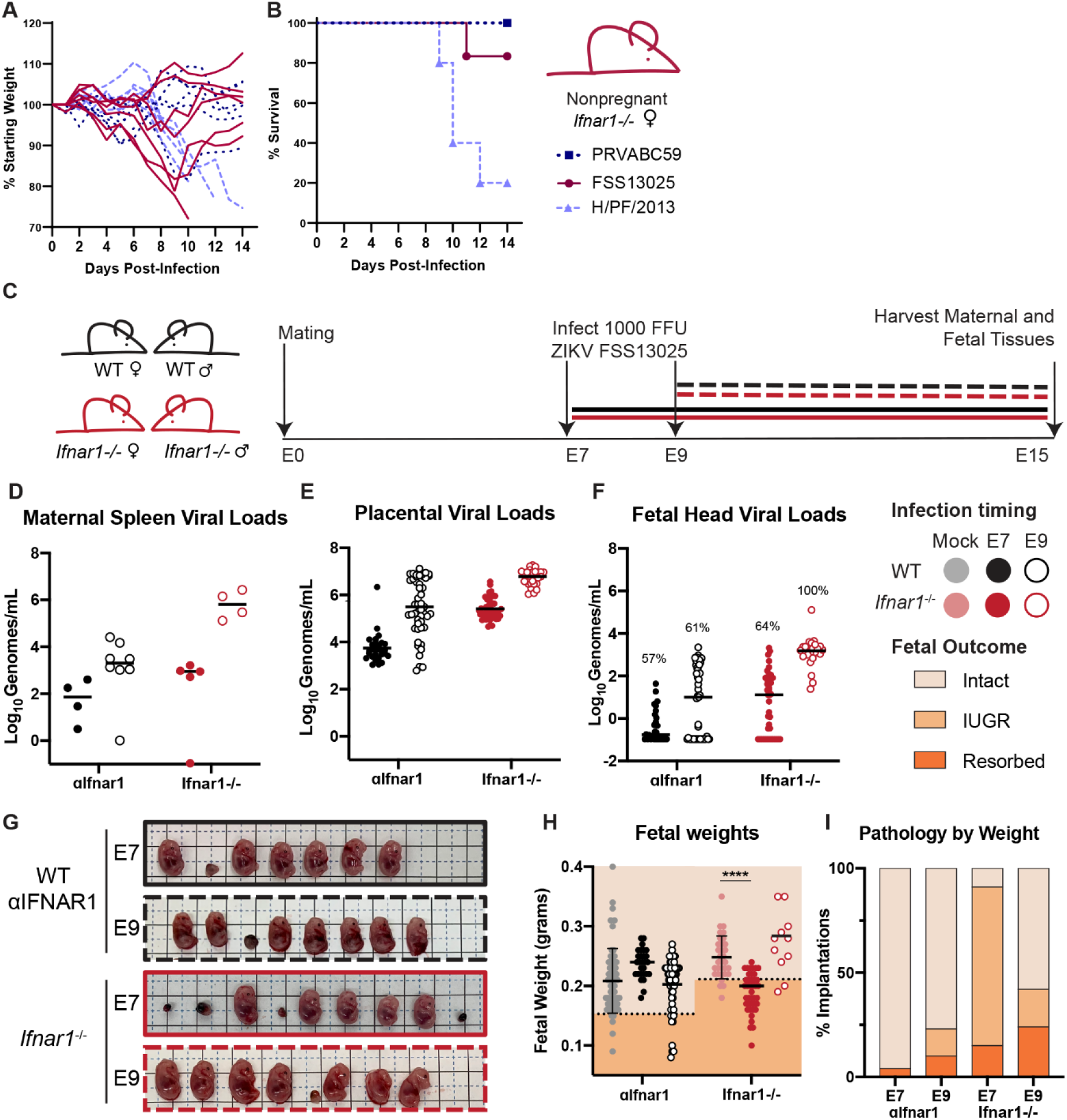
Infection earlier in gestation corresponds to enhanced fetal pathology in mouse models of congenital ZIKV infection. **A-B.** Non-pregnant 8-10 week old female *Ifnar1*^-/-^ mice (5-6 mice per group) were infected with 1000 FFU of ZIKV strain PRVABC59, FSS13025, or H/PF/2013; weights and survival were measured daily for 14 days. Each line represents an individual mouse. **C-K.** Dams from WT x WT or *Ifnar1*^-/-^ x *Ifnar1*^-/-^ crosses (6 to 8 WT or 4 to 5 *Ifnar1*^-/-^ dams per group) were infected at day 7 or 9 post-mating (E7, E9) with ZIKV FSS13025 by subcutaneous inoculation in the footpad. WT dams were given 2mg of anti-IFNAR1 blocking mAb intraperitoneally one day prior to infection. Tissues were harvested at E15 (8 or 6dpi). **D-F**. ZIKV viral loads in the maternal spleen, placenta, and fetal head were measured by qRT-PCR. Each data point represents one dam (**D**) or fetus (**E-F**). The percent of ZIKV-positive fetal heads is indicated above each group. **G**. Representative images of fetuses/resorptions from one pregnancy from each cross. **H.** Intact fetuses (i.e. not resorbed) were weighed. Fetuses <1 standard deviation from the mean of mock-infected (below dotted line) were classified as having intrauterine growth restriction (IUGR). Intact fetuses with weights significantly different from mock pregnancies (calculated by ANOVA) are indicated, **** P<0.0001. **I**. Proportions of fetuses exhibiting IUGR or resorption.

To determine if we could observe similar pregnancy pathology with another virus that causes congenital infections in humans, we sought to generate a RUBV mouse model, as small animal models to study RUBV pathogenesis are not available and experimental RUBV infections in knockout mice have not been reported. We first infected 8-week-old non-pregnant wild-type, *Ifnar1^-/-^*, and *Ifnlr1*^-/-^ mice with 1000 FFU and 5-week-old *Ifnar1*^-/-^ mice with 1x10^5^ FFU of RUBV (strain M33) by intranasal inoculation or subcutaneous inoculation in the footpad but observed no weight loss or disease signs (Figure S2A-B). To determine whether mice supported any RUBV infection, we inoculated *Ifnar1^-/-^ Ifngr1^-/-^* DKO mice intravenously with 1x10^5^ FFU of RUBV and measured viral RNA by qRT-PCR from blood and serum at 2, 4, and 7 dpi and from spleen, lung, and kidney at 7 dpi (Figure S2C-D). Although in humans RUBV targets a variety of tissues and produces viremia (30), we found very low or undetectable viral loads in *Ifnar1^-/-^ Ifngr1^-/-^* DKO mice even though these mice are highly susceptible to many viral infections. Since human congenital rubella syndrome requires maternal viremia, we concluded that this mouse model would not be suitable for assessing transplacental transmission of RUBV and limited our further studies to ZIKV.

### Mid-gestation mouse placentas produce IFN-λ in the presence and absence of infection

IFN-λ is secreted constitutively from human primary trophoblasts cultured *ex vivo*, trophoblast organoids, and placental cell lines grown in 3D culture (6–8). Although IFN-λ has antiviral activity at the murine maternal-fetal interface (9), it was unknown if IFN-λ was secreted constitutively from the mouse placenta. To evaluate IFN-λ activity in the absence of infection, we measured IFN-λ activity from placentas harvested from mid-to-late gestation (E11 to labor). In uninfected mice we found that placental IFN-λ activity varied considerably over the course of gestation, increasing from E11 to E15 then dropping at E17 (Figure 2A). IFN-λ activity rose again in placentas taken from dams in active labor, consistent with the cytokine response that triggers parturition. We also detected IFN-λ in the placentas of ZIKV-infected dams harvested at E15 (Figure 2B). Placentas from MAR1-5A3-treated WT dams infected at E7 had significantly higher IFN-λ activity than placentas from dams infected at E9, but there was no effect of infection timing in *Ifnar1*^-/-^ dams. Unexpectedly, IFN-λ activity in placentas from ZIKV-infected dams was reduced compared to uninfected WT dams harvested at E15. These results indicate that IFN-λ is constitutively expressed during mouse pregnancy, and also is present during congenital ZIKV infection.

**Figure 2.**
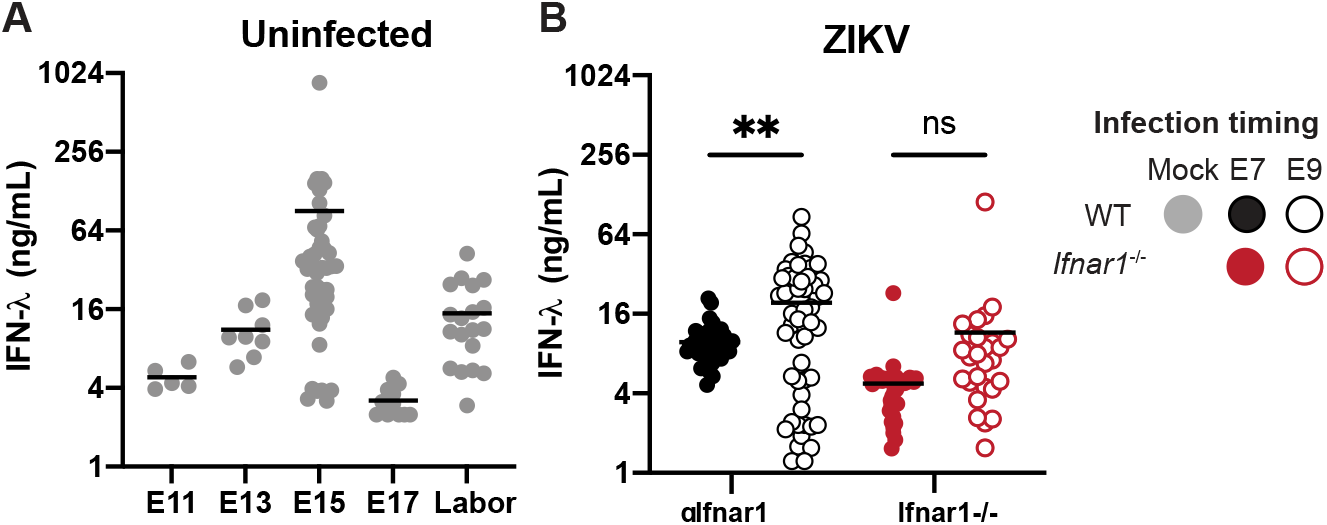
IFN-λ is produced at the maternal-fetal interface. **A.** Placentas were harvested from uninfected pregnant dams at E11 (1 dam), E13 (1 dams), E15 (5 dams), E17 (3 dams), and during labor (3 dams). Placentas were homogenized in PBS, and IFN-λ activity in placental homogenate was determined using a reporter cell line. **B.** Pregnant WT dams (treated with an anti-IFNAR1 blocking mAb, 6 or 8 dams per group) or pregnant *Ifnar1*^-/-^ dams (4 or 5 dams per group) were infected at either E7 or E9 with ZIKV FSS13025. Placentas were homogenized in PBS, and IFN- λ activity in placental homogenate was determined using a reporter cell line.

### Maternal IFN-λ signaling restricts ZIKV transplacental transmission in a gestational stage-dependent manner

Prior studies found that IFN-λ signaling reduces ZIKV transplacental transmission in mice (9) but the cells and tissues responding to IFN-λ were not identified. To determine the targets of IFN-λ signaling at the maternal-fetal interface we first assessed whether the protective effects of IFN-λ were mediated by signaling in maternal or fetal tissues. To generate pregnancies with distinct maternal and fetal IFN-λ responsiveness, we crossed *Ifnlr1*^+/-^ dams by *Ifnlr1*^-/-^ sires, or the reverse, producing litters comprising *Ifnlr1^+/-^* and *Ifnlr1^-/-^* fetuses within dams that either retained IFN-λ signaling (*Ifnlr1^+/-^)* or lacked it (*Ifnlr1^-/-^)* (Figure 3A). We infected mice with 1000 FFU of ZIKV FSS13025 by subcutaneous inoculation in the footpad at E9, 1 day following administration of 2mg of MAR1-5A3. At 6 dpi (E15), we harvested maternal and fetal tissues and determined fetal *Ifnlr1* genotype by PCR, and viral loads by qRT-PCR. We found no difference in maternal or placental viral loads based on maternal or fetal *Ifnlr1* genotype (Figure 3B-C). In contrast, we found higher rates of ZIKV transplacental transmission in dams lacking IFN-λ signaling (*Ifnlr1^-/-^)*, regardless of fetal genotype (67% vs 28%) and viral loads were significantly higher in fetuses from in *Ifnlr1*^-/-^ dams compared to *Ifnlr1*^+/-^ dams, regardless of fetal genotype (P<.0001) (Figure 3D). Higher viral loads were not accompanied by overt pathology in this model as there was no difference in fetal weights (Figure 3E) based on either maternal or fetal *Ifnlr1* genotype. Our observation of higher viral loads in the fetuses of *Ifnlr1*^-/-^ dams, regardless of fetal *Ifnlr1* genotype, provides strong evidence that IFN-λ signaling protects against transplacental transmission of ZIKV via signaling exclusively in maternal tissues. This is specific to tissues at the maternal fetal-interface, as non-pregnant female *Ifnlr1^-/-^* and *Ifnlr1^+/-^* mice exhibited no differences in viremia or tissue viral loads following infection (Figure 3F-G).

**Figure 3.**
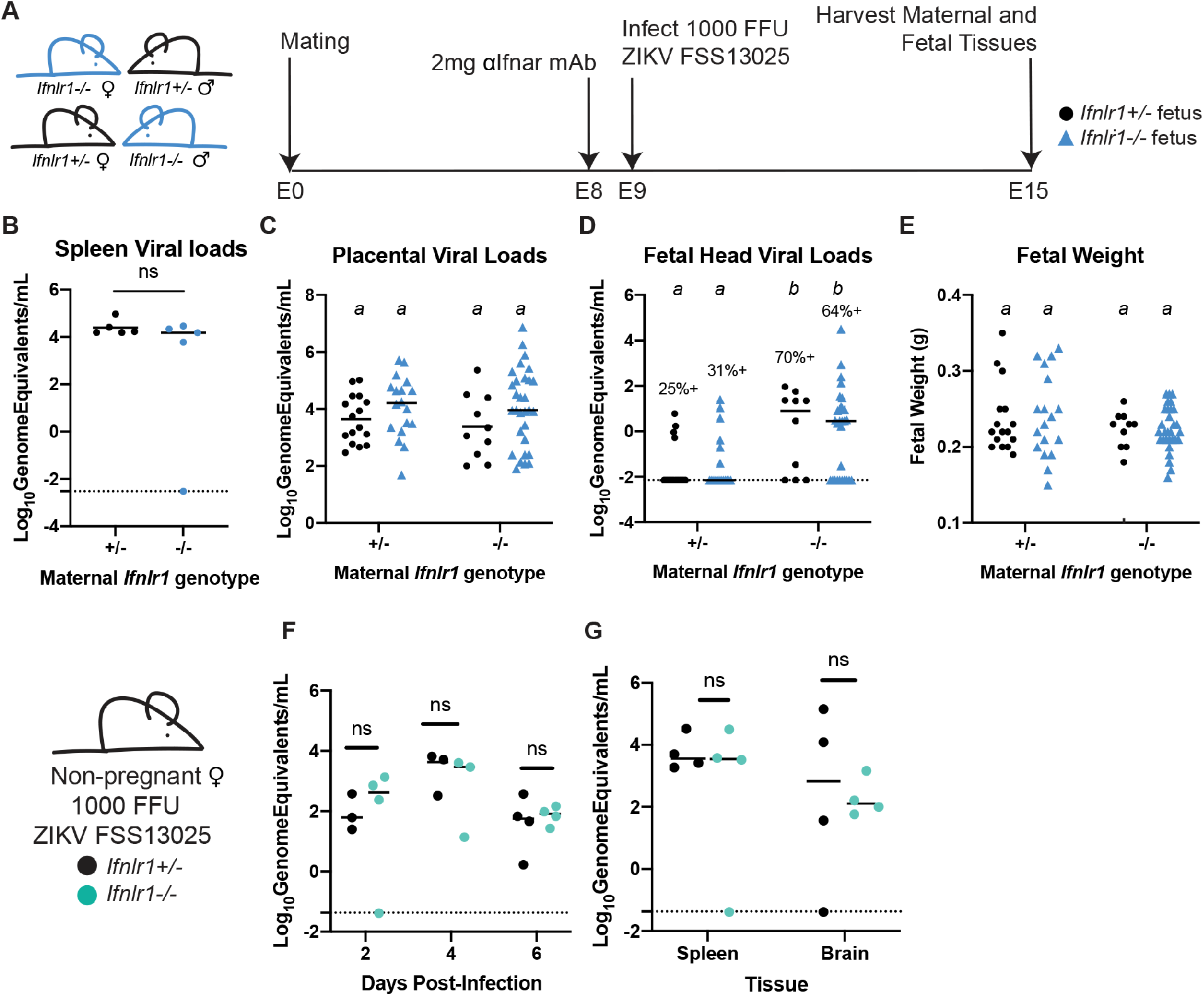
IFN-λ restricts ZIKV transplacental transmission by signaling to maternal tissues. **A**. Mating and infection timeline. *Ifnlr1*^+/-^ dam x *Ifnlr1*^-/-^ sire and *Ifnlr1*^-/-^ dam x *Ifnlr1*^+/-^ sire crosses were used to generate pregnancies with IFN-λ responsive (*Ifnlr1*^+/-^) and non-responsive (*Ifnlr1*^-/-^) fetuses. **B-E.** Pregnant dams were treated with 2mg of IFNAR1 blocking antibody at E8 and infected at E9 with 1000 FFU of ZIKV FSS13025 by subcutaneous inoculation in the footpad. Fetuses and their associated placentas were harvested at E15. ZIKV RNA was measured by qRT-PCR in maternal spleen (**B**), placenta (**C**), and fetal head (**D**) and fetuses were weighed (**E**). The percent of fetuses with detectable ZIKV is noted (**D**). Data are combined from 5 or 6 dams per group; each data point represents a single dam (**B**) or fetus (**C-E**). Groups were compared by ANOVA (**B, C, E**) or Mann-Whitney (**D**); italicized letters indicate groups that are significantly different each other (P < 0.05). **F** and **G.** Non-pregnant, 8-week old *Ifnlr1*^-/-^ and *Ifnlr1*^+/-^ females were infected with 1000 FFU of ZIKV FSS13025. Viremia was measured from serum at 2, 4, and 6 dpi by qRT-PCR. Spleens and brains were harvested 6 dpi, and viral loads were measured by qRT-PCR. Groups were not significantly different (ns) by ANOVA.

In mice placental differentiation is complete around E10.5 (31) so mice infected at E9 are expected to have a fully-formed placenta by the time ZIKV reaches the placenta from maternal circulation. Since pregnancy pathology depends on gestational stage at the time of infection (Figure 1) (9, 29), and IFN-λ antiviral effects vary with gestational time (Jagger et al., 2017) we next assessed the effects of IFN-λ signaling in maternal and fetal tissues following ZIKV infection two days earlier, at E7. At this earlier infection time, maternal viremia is expected to be established prior to complete placentation. We again crossed *Ifnlr1*^+/-^ and *Ifnlr1*^-/-^ mice to generate pregnancies with mixed IFN-λ responsiveness within dams that could or could not respond to IFN-λ (Figure 4A). We infected pregnant dams with 1000 FFU of ZIKV FSS13025 by subcutaneous inoculation in the footpad at E7, 1 day following administration of 2mg of MAR1-5A3. At 8 dpi (E15), we harvested maternal and fetal tissues and determined *Ifnlr1* genotype by PCR and viral loads by qRT-PCR. Similar to infection at E9, we found no difference in maternal spleen or placental viral load in *Ifnlr1*^-/-^ compared to *Ifnlr1*^+/-^ dams (Figure 4B-C). However, in contrast to the protective effect of maternal IFN-λ signaling after E9 infection, with E7 infection we found higher rates of transplacental transmission in *Ifnlr1*^+/-^ dams compared to *Ifnlr1*^-/-^ (47% versus 13%, *P* =.0006) (Figure 4D). Moreover, fetuses from *Ifnlr1*^+/-^ dams had significantly higher viral burdens than those from *Ifnlr1*^-/-^ dams, regardless of fetal genotype (Figure 4D). Although we found significant differences in fetal weights from *Ifnlr1*^+/-^ and *Ifnlr1*^-/-^ dams (*P* < 0.0001), the difference results from an increase in fetal weights from *Ifnlr1*^+/-^ pregnancies and *Ifnlr1^-/-^* fetal weights fell within the range of uninfected pregnancies (Figure 1 I-J). Altogether these results show that IFN-λ signaling exerts a gestational stage-specific effect on ZIKV transplacental transmission, where earlier in gestation IFN-λ signaling facilitates ZIKV transplacental transmission in contrast to later stages where IFN- λ inhibits transplacental transmission. Importantly, at either stage, the effects of IFN-λ signaling were mediated through signaling in maternal tissues, rather than through signaling in the placenta or fetus, as only maternal *Ifnlr1* genotype influenced ZIKV transmission, not fetal *Ifnlr1* genotype.

**Figure 4.**
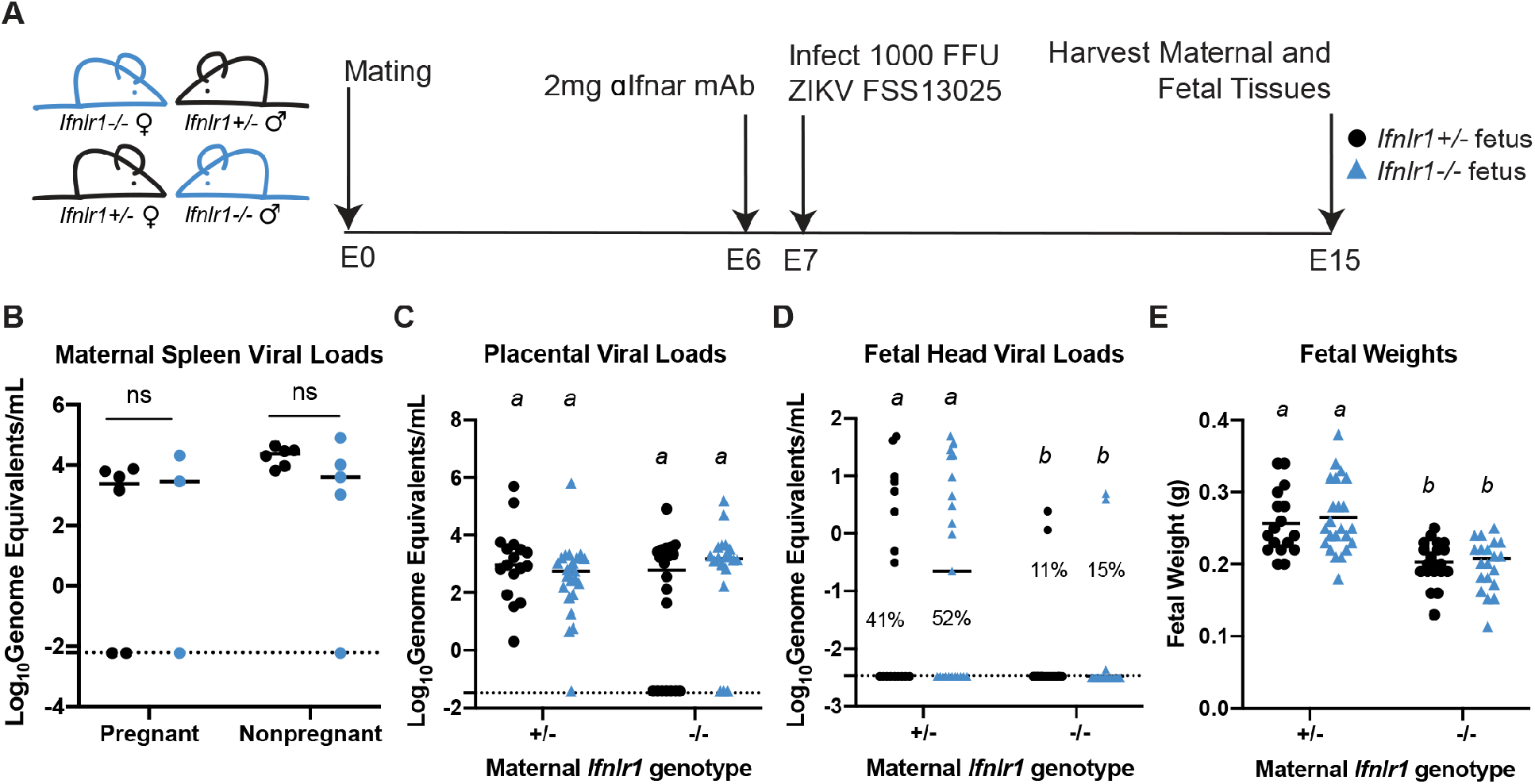
IFN-λ enhances fetal infection early in gestation through signaling to maternal tissues. **A**. Mating and infection timeline. *Ifnlr1*^+/-^ dam x *Ifnlr1*^-/-^ sire and *Ifnlr1*^-/-^ dam x *Ifnlr1*^+/-^ sire crosses were treated with 2mg of anti-IFNAR1 mAb at E6 and infected at E7 with 1000 FFU of ZIKV FSS13025 by subcutaneous inoculation in the footpad. Fetuses and their associated placentas were harvested at E15. **B-D.** ZIKV RNA in the maternal spleens, placenta, and fetal head were measured by qRT-PCR. **E.** Gross fetal pathology was measured by fetal weight. Significant differences are denoted by italicized letters, calculated by ANOVA (**B, C, E**) or Mann- Whitney (**D**). Data are combined from 5 or 6 dams per group; each data point represents a single dam (**B**) or fetus (**C-E**).

### Maternal IFN-λ signaling exacerbates fetal pathology early in gestation

Since maternal IFN-λ signaling enhanced rather than limited ZIKV transmission at E7, we next assessed the effects of IFN-λ signaling on fetal pathology at this early gestational stage. These experiments used mice that retained or lacked IFN-λ signaling on an *Ifnar1*^-/-^ background, as we did not observe overt pathology in IFNAR1-intact mice (Figure 1G-J). We crossed wild-type, *Ifnar1^-/-^,* and *Ifnar1^-/-^Ifnlr1^-/-^* dams and sires to generate pregnancies in which IFN-λ signaling was present or absent on both or either side of the maternal-fetal interface (Figure 5A). Pregnant dams were infected with ZIKV FSS13025 at E7, and tissues were harvested 8 dpi (E15). Pregnancies that lacked both IFN-αβ and IFN-λ signaling on both sides of the maternal-fetal interface exhibited significant growth restriction compared to uninfected pregnancies (Figure 5B-D, group 1). Fetal IFN-αβ signaling previously has been shown to be pathogenic during congenital ZIKV infection in mice (29) and accordingly we found that all fetuses were resorbed in pregnancies that retained IFN-αβ and IFN-λ signaling exclusively on the fetal side of the interface (Figure 5B-D, group 2). This pathology was mediated by fetal IFN-αβ signaling because when fetal IFN-λ signaling was restored in the absence of fetal IFN-αβ signaling, we found no resorptions (Figure 5B-D, group 3). In contrast, when IFN-λ signaling was restored on both the fetal and maternal side, 30% of the fetuses were resorbed and the remaining intact fetuses were significantly smaller than those from uninfected pregnancies (Figure 5B-D, group 4). Moreover, pregnancies with maternal IFN-λ signaling had variable fetal outcomes (Figure 5D), both within and between pregnancies (Figure 5E). There were no differences in maternal spleen viral loads or transplacental transmission as determined by qRT-PCR (Figure 5F-G). Since pregnancies with maternal IFN-λ signaling exhibited variable pathologic outcomes within litters, we asked whether this was influenced by fetal sex. We determined fetal sex by PCR genotyping for *Sry,* a gene found on the Y chromosome, and found that 40% of male fetuses were resorbed (20% of total implantations) compared to 6% of female fetuses (3% of implantations) (Figure 5H). This raises the possibility that IFN-λ mediated outcomes could be driven by maternal immune rejection, as only male fetuses are genetically distinct from the mother in congenic mouse pregnancies. Since our results and prior studies (29) showed that IFN signaling can be pathogenic in the context of congenital ZIKV infection, we considered whether IFN signaling might be detrimental during pregnancy more generally. However, in analyzing ∼17 months of breeding records from WT, *Ifnar1*^-/-^, *Ifnlr1*^-/-^, and *Ifnar1*^-/-^ *Ifnlr1*^-/-^ mice in our colony (>275 litters from >40 breeder cages) we found no significant difference in litter size between the lines, supporting the idea that IFN signaling during pregnancy is not detrimental outside an infection or other inflammatory context. We found no difference in viremia or tissue viral loads between *Ifnar1*^-/-^ and *Ifnar1*^-/-^ *Ifnlr1*^-/-^ non- pregnant females (Figure 5I-J), altogether indicating that the pathogenic effects of IFN-λ at the maternal-fetal interface are distinct from restricting viral replication systemically in the dam.

**Figure 5.**
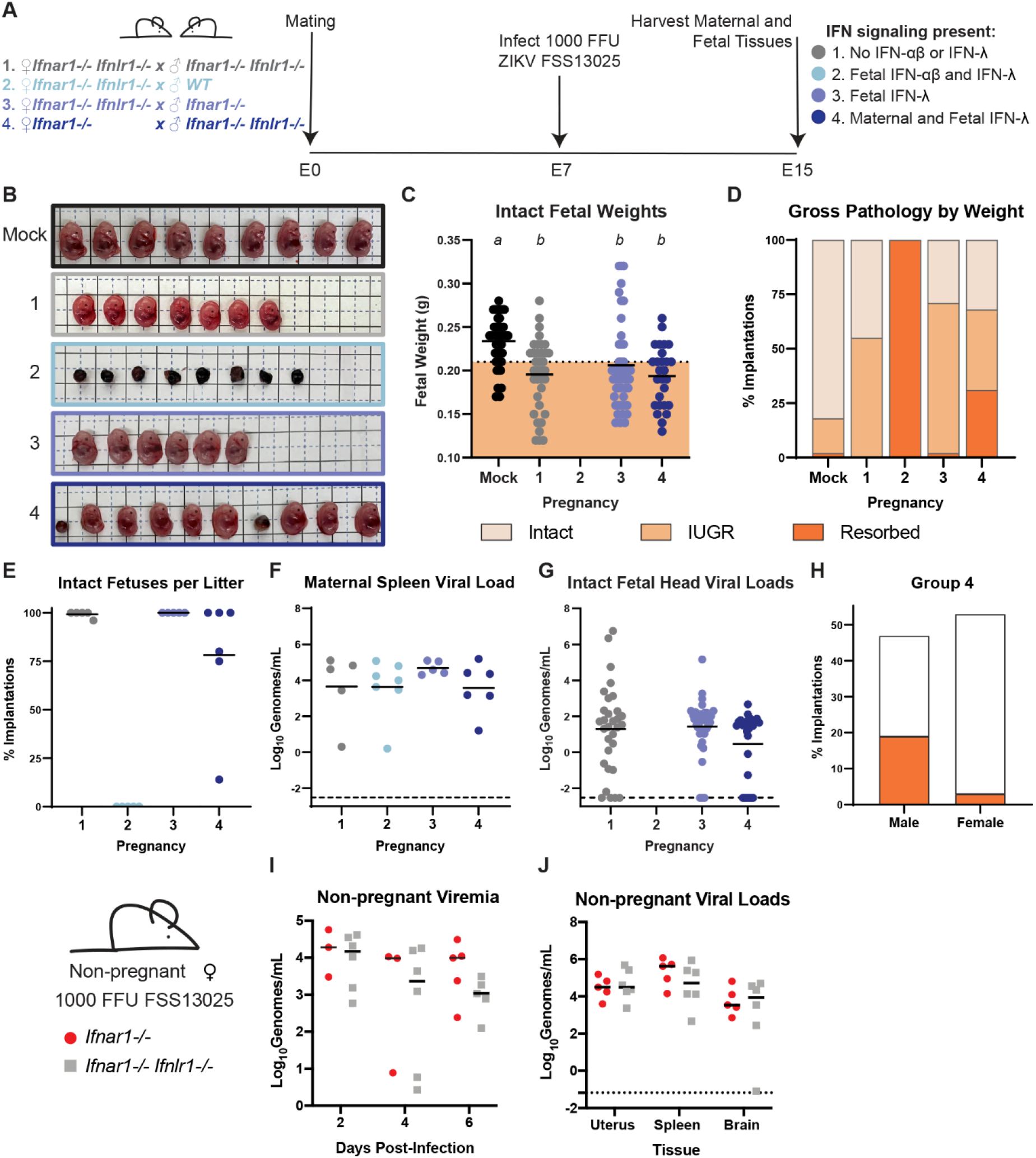
Maternal IFN-λ signaling induces fetal pathology. **A.** Mating and infection timeline. Wild-type, *Ifnar1^-/-^, and Ifnar1^-/-^ Ifnlr1*^-/-^ DKO mice were crossed to create pregnancies with differing IFN-λ responsiveness in maternal and fetal tissues, within dams lacking IFN-αβ signaling. Pregnant dams were infected at E7 with 1000 FFU of ZIKV FSS13025 by subcutaneous inoculation in the footpad. Data are combined from 5 to 7 dams per group. **B**. Representative images of the fetuses/resorptions from each cross. **C.** Intact fetuses (not resorbed) were weighed. Fetuses with weights below one standard deviation of uninfected pregnancies were classified as having IUGR. Significant differences between fetal groups are indicated by italicized letters and were calculated by ANOVA. **D**. The percent of resorptions and IUGR in each pregnancy group. **E.** The percent of intact fetuses in individual litters. **F** and **G**. ZIKV viral loads in fetal head and maternal spleen were measured by qRT-PCR. **H.** The sex of resorptions and intact fetuses was determined by PCR. **I** and **J**. 10-week-old non-pregnant females were infected with 1000 FFU of ZIKV FSS13025 by subcutaneous inoculation in the footpad. Viral loads in serum (2, 4, 6 dpi) and tissues (6 dpi) were determined by qRT-PCR.

### IFN-λ pathogenic effects are mediated by leukocytes and decrease over gestational time

To determine whether IFN-λ-mediated fetal pathology was specific to ZIKV infection, we assessed the pathogenic effect of IFN-λ signaling stimulated by poly(I:C). To determine which tissues produced IFN in response to poly(I:C) treatment, we measured IFN-λ and IFN-β in serum, uterus, lung, and spleen 24 hours post poly(I:C) treatment in pregnant and non-pregnant WT mice. We detected IFN-λ activity only in uteruses from pregnant mice (Figure 6A), while IFN-β was detected in tissues but not serum (Figure 6B), confirming that poly(I:C) treatment can induce IFN-λ and IFN-αβ at the maternal-fetal interface. We next assessed the effect of poly(I:C)-treatment on fetal pathology in *Ifnlr1*^+/-^ and *Ifnlr1*^-/-^ dams mated to WT sires. To investigate the possibility that IFN-λ mediated pathology resulted from maternal immune signaling, we also included dams lacking IFN-λ signaling in hemopoietic cells *(Vav*-Cre *Ifnlr1^-/-^,* Figure S3) mated to WT sires. We administered 200 µg of poly(I:C) by intraperitoneal injection to dams at E7, E9, or E11 and assessed fetal outcome at E15 (Figure 6C). Consistent with our observations in ZIKV-infected *Ifnar1^-/-^* dams, *Ifnlr1*^+/-^ dams exhibited a 7.5-fold higher resorption rate compared to *Ifnlr1*^-/-^ dams after poly(I:C) administration at E7 (31% vs 3%, Figure 6D-E). *Vav*-Cre *Ifnlr1^-/-^* dams also had low rates of fetal resorption (3%), indicating that IFN-λ mediated pregnancy pathology acts through maternal immune cells. Accordingly, we found that decidualized human endometrial cells supported replication of ZIKV and RUBV and responded to IFN-β treatment but did not respond to IFN-λ treatment (Figure S4). Although IFN-λ signaling induced resorptions, the weights of intact fetuses were no different in poly(I:C)-treated dams compared to mock-treated, indicating that poly(I:C) treatment does not induce an IUGR phenotype (Figure 6F). We also observed IFN-λ induced resorptions following poly(I:C) administration at E9 and E11 (Figure 6E), but the effect was less pronounced compared to treatment at E7, consistent with a model where the pathogenic effects of maternal IFN-λ signaling are most severe earlier in gestation.

**Figure 6.**
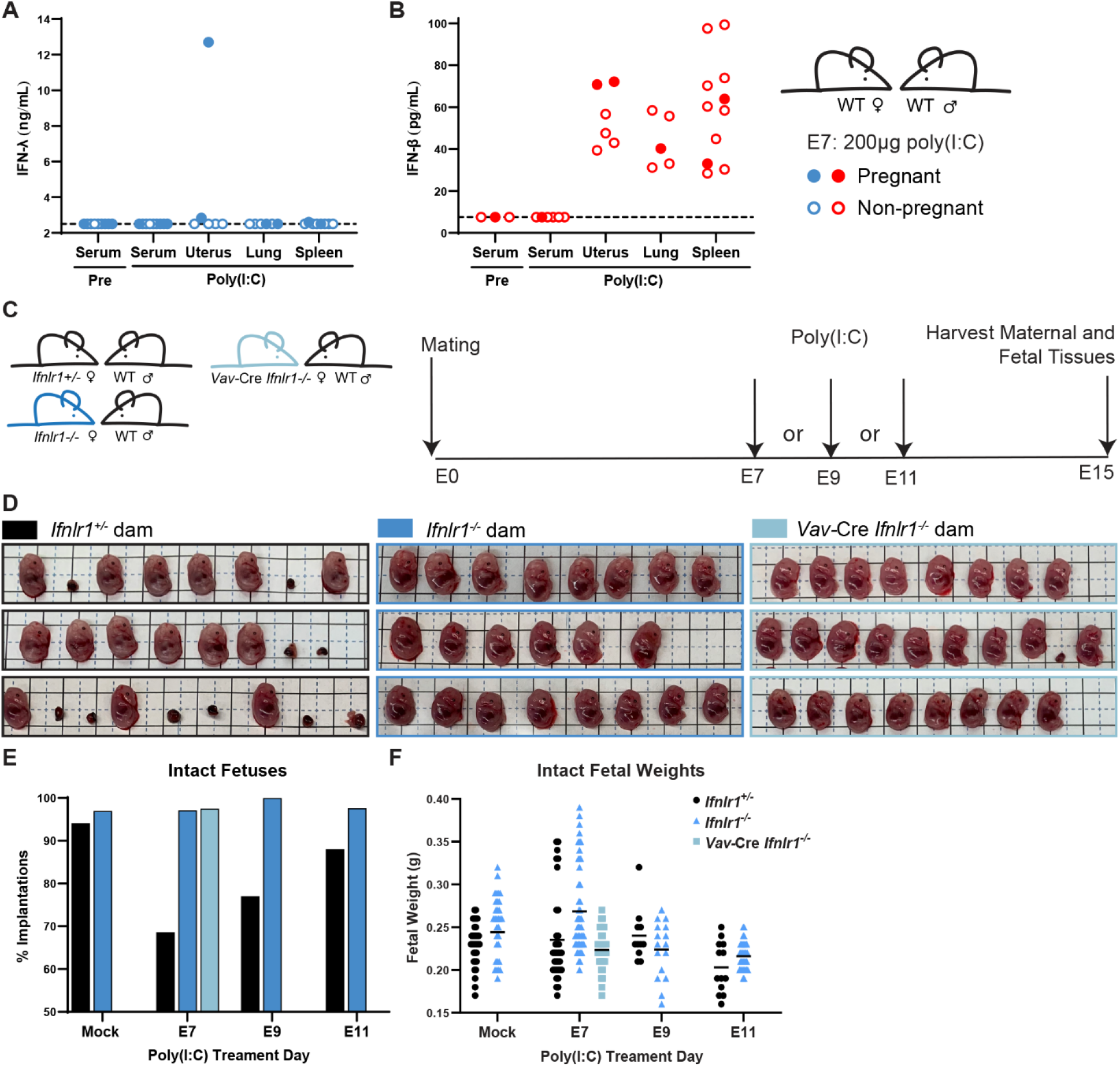
IFN-λ mediates fetal pathology through signaling in maternal leukocytes. A and B. WT dams were mated to WT sires and treated with 200 µg of poly(I:C) at E7. Serum was collected by submandibular bleed pre-treatment. 24 hours post-treatment, the uterus, spleen, and lung were harvested from pregnant and non-pregnant mice. IFN-λ activity was measured in a reporter cell assay and IFN-β concentration by ELISA. Filled circles represent pregnant mice. **C**. Experiment timeline. *Ifnlr1^+/-^, Ifnlr1^-/-^,* and leukocyte *Ifnlr1^-/-^* (*Vav*-Cre *Ifnlr1^-/-^*) dams were mated to WT sires. Pregnant dams were administered 200µg of poly(I:C) by intraperitoneal injection at E7, E9, or E11 and fetuses were harvested at E15. **D.** Representative litters from E7-treated pregnancies. Resorptions were counted **(E)** and intact fetuses were weighed **(F).**

## DISCUSSION

Our results show that IFN-λ can have both protective and pathogenic effects during pregnancy depending on gestational stage, but that both effects occur via signaling in maternal tissues. This identifies a distinct role for IFN-λ compared to IFN-αβ at the maternal-fetal interface as the pathologic effects of IFN-αβ act through signaling in fetal tissues in similar ZIKV congenital infection models (29). The contrasting effects of maternal IFN-λ signaling at different gestational stages likely derive from differences in the physiology of the maternal-fetal interface over the course of gestation, producing distinct outcomes when ZIKV infects the maternal-fetal interface prior to placentation (maternal inoculation at E7) or after the placenta has formed (E9 inoculation). We further showed that IFN-λ mediated pathology is mediated by leukocytes after poly(I:C) treatment. Because the maternal immune landscape varies over gestation, IFN-λ may signal to a leukocyte population that diminishes or changes as pregnancy progresses.

Since IFN-λ is constitutively secreted from human trophoblasts and these cells are refractory to replication by a wide array of viruses and other infectious agents (32–34), we had expected IFN-λ to restrict ZIKV transplacental transmission by signaling on placental trophoblasts and inducing a cell-intrinsic antiviral response. Instead, we found that IFN-λ antiviral activity was mediated through signaling in maternal tissues. Importantly, pregnant and non-pregnant *Ifnlr1*^-/-^ mice showed no differences in viral loads in peripheral tissues (serum, spleen), consistent with prior studies with other flaviviruses (35, 36) and excluding that enhanced ZIKV transplacental transmission is due to enhanced viral replication and spread in maternal tissues. The mechanism by which maternal IFN-λ signaling restricts ZIKV transplacental transmission remains unclear, but could include antiviral activity in the uterine decidua, or immunomodulatory effects on maternal leukocytes, such as decidual NK cells, Tregs, neutrophils or dendritic cells. We did not observe IFN-λ responsiveness in a human decidualized endometrial cell line, but decidual cells respond to IFN-λ in other human cell culture models, including explants and organoids (8, 9). Differences in decidual responsiveness have been noted from cell models taken at different times in gestation (37) and could explain the lack of IFN-λ responsiveness we observed in a decidual cell line.

Although we did not find a role for fetal IFN-λ signaling, human placental models do respond to IFN-λ in culture. Differences in placental IFN-λ responsiveness could be due to variations in the mouse and human placentas. Although both are discoid and hemochorial, mouse and human placentas have distinct trophoblast lineages which include trophoblast giant cells and a second layer of syncytiotrophoblasts in mice and extravillous trophoblasts in humans (31).

In contrast to the protective effect of IFN-λ that we observed later in gestation, IFN-λ signaling enhanced transplacental transmission in mice infected at E7. We attribute this enhanced transmission to pathogenic effects of IFN-λ signaling on the placental barrier which is not yet fully formed at this stage of gestation. Remarkably, these pathogenic effects were mediated exclusively by IFN-λ signaling in maternal tissues, similar to the protective effects of IFN-λ. Interestingly, the pathogenic effects of IFN-λ did not require ZIKV infection, as we could elicit a similar phenotype by treating pregnant dams with poly(I:C). We found that rates of IFN-λ mediated pathology decreased when poly(I:C) was administered later in gestation. Congenital viral infections also produce fetal pathology with gestational stage-dependent effects in humans: congenital rubella syndrome almost entirely results from infection in the first trimester and ZIKV and human cytomegalovirus (HCMV) infections early in pregnancy likewise produce the most severe outcomes, although ZIKV and HCMV can be deleterious throughout pregnancy (1). Our findings emphasize the importance of studying congenital infections and immune responses at different gestational stages. One limitation of this study is that it does not include infections at time points following placentation, so it remains to be determined how IFN-λ affects pregnancies late in gestation. Furthermore, congenital infections can result from both transplacental spread and ascending infection from the vagina. The immunologic and anatomical barriers to ascending infection are different from those to transplacental infection, so IFN-λ could have distinct effect based on the route of infection.

We found that fetuses were protected from resorption when IFN-λ signaling was ablated only in hematopoietic cells, indicating that IFN-λ pathogenic effects result from signaling in maternal leukocytes. Since IFN-λ mediated pathology depends on gestational stage, IFN-λ may act through particular leukocytes present earlier in gestation that diminish over time. Early in gestation, 40% of the maternal decidua is made up of leukocytes including NK cells, macrophages, and Tregs (2). These populations change over the course of gestation and play a critical role in mediating placental invasion and spiral artery formation. IFN-λ signals to several of these cell types in contexts outside of pregnancy (38) and potentially could disturb the immune balance necessary for proper placentation. Multiple distinct subsets of macrophages have been identified in the maternal decidua, and imbalance between macrophages subtypes is associated with adverse pregnancy outcomes (39, 40). IFN-λ changes the transcriptional profile and increases pro-inflammatory phenotypes of monocytes differentiated into macrophages in culture (41, 42). Macrophages skew towards a M2 phenotype as pregnancy progresses and IFN-λ could increase proportions of M1 macrophages, potentially leading to inflammation and fetal rejection. Placentas harvested from ZIKV-infected rhesus macaques have more monocytes and macrophages than those from uninfected animals, as well as changes in the proportions of monocyte subsets (43). Although a function for neutrophils at the maternal-fetal interface has not been well defined, mouse neutrophils do respond to IFN-λ. However, IFN-λ has anti-inflammatory activity in these contexts and is associated with reductions in inflammatory pathology during influenza infection as well as rheumatoid arthritis (44, 45). Further research focusing on identifying the specific maternal cell types that respond to IFN-λ signaling will enhance our understanding of the mechanisms underlying IFN-λ-mediated fetal pathology.

We found a striking sex difference in IFN-λ mediated pathology, with male fetuses exhibiting significantly higher resorption rates than female fetuses. This observation is consistent with immune-mediated rejection, as only male fetuses are genetically distinct from the dam in these congenic pregnancies. Immunity at the maternal-fetal interface is carefully regulated to prevent non-self-rejection of the fetus, and includes mechanisms that downregulate NK cell cytotoxicity and recognition of non-self-tissues (2). Modeling congenital infection in semi-allogeneic pregnancies will provide further insight into the role of IFN-λ signaling in changes to maternal immune tolerance.

Although IFN-λ is best-characterized for its protective activity in the context of viral infections, particularly in the respiratory and gastrointestinal tracts (5), IFN-λ signaling also is associated with deleterious effects in some other contexts. IFN-λ contributes to impaired tissue repair following respiratory and gastrointestinal infections in mice (46–48). In humans, multiple polymorphisms in the IFN-λ locus are associated with clinical outcomes from hepatitis C virus (HCV) infection (49). Among these, a frameshift mutation in the promoter of *IFNL4* results in the loss of IFN-λ4 production and concomitant improved clearance of HCV as well as other gastrointestinal and respiratory infections though the mechanism by which the loss of an IFN results in an improved antiviral response remains unclear (50, 51). The pseudogenization of IFN- λ4, along with selection for lower-potency variants, suggest IFN-λ4 signaling has been deleterious during human evolution (52, 53). In mice, the IFN-λ family consists only of IFN-λ2 and IFN-λ3 as IFN-λ1 is a pseudogene and the genomic region encoding IFN-λ4 is absent (54, 55), which limits some comparisons of the effects of IFN-λ in mice and humans.

Our observations of a pathogenic effect of IFN-λ signaling at the maternal-fetal interface bear some similarity to the pathogenic effects of IFN-αβ in pregnancy, though notably in mouse models of congenital ZIKV infection IFN-αβ is pathogenic when it signals to fetal tissues (29) whereas we find that IFN-λ acts through signaling in maternal tissues. Women with dysregulated IFN-αβ signaling (sustained IFN production or impaired receptor downregulation) exhibit poor pregnancy outcomes including pre-eclampsia as well as neurodevelopmental defects similar to those induced by congenital infection, consistent with a role for dysregulated IFN-αβ responses in placental damage (56–60). Whether dysregulated IFN-λ signaling exerts similar effects during human pregnancy remains to be determined.

Altogether, these findings identify an unexpected effect of IFN-λ signaling specifically in maternal (rather than placental or fetal) tissues, which is distinct from the pathogenic effects of IFN-αβ during pregnancy. These results highlight the complexity of immune signaling at the maternal-fetal interface, where disparate outcomes can result from signaling at different gestational stages.

## MATERIALS AND METHODS

### Viruses

Virus stocks were grown in Vero cells in Dulbecco’s modified Eagle medium (DMEM) containing 5% fetal bovine serum (FBS), L-glutamine, and HEPES at 37°C with 5% CO2. ZIKV strain FSS13025 (Cambodia 2010) was obtained from the World Reference Center for Emerging Viruses and Arboviruses (61). ZIKV strains PRVABC59 (Puerto Rico 2015) and H/PF/2013 (French Polynesia 2013) were obtained from U.S. CDC (62, 63). Rubella virus strain M33 was obtained from Dr. Michael Rossmann, Purdue University (64). DENV4 (TVP-360) was obtained from Dr. Aravinda DeSilva, UNC Chapel Hill. Virus stock titer was quantified by focus-forming assay on Vero cells (65). Viral foci were detected using 500 ng/mL of anti-flavivirus mouse monoclonal antibody E60 (66) or 1:1000 dilution of goat anti-RUBV antibody (Lifespan Biosciences LC-C103273/39321), 1:5000 dilution of an HRP conjugated goat anti-mouse IgG (Sigma #A8924) or 1:5000 dilution of an HRP conjugated rabbit anti-goat (Sigma #A5420), and

TrueBlue peroxidase substrate (KPL). Antibody incubations were performed overnight at 4°C. Foci were counted on a CTL Immunospot analyzer.

### Mice

All experiments and husbandry were performed under the approval of the University of North Carolina at Chapel Hill Institutional Animal Care and Use Committee. Experiments used 8–20- week-old female mice on a C57BL/6 background. Wild-type mice were obtained commercially (Jackson labs 000664) or bred in-house. *Ifnar1*^-/-^ and *Ifnar1^-/-^Ifngr1^-/-^* mice were obtained from Dr. Jason Whitmire (UNC) then bred in-house. *Ifnlr1*^-/-^ mice were provided by Dr. Herbert Virgin (Washington University in St. Louis), generated by crossing *Ifnlr1*^fl/fl^ mice with mice constitutively expressing Cre recombinase under a CMV promoter (67); these mice were then bred in-house as knockout x knockout (36);. *Ifnar1*^-/-^ *Ifnlr1*^-/-^ DKO mice were generated by crossing *Ifnlr1*^-/-^ and *Ifnar1*^-/-^ mice. *Ifnlr1*^+/-^ mice were generated by crossing *Ifnlr1*^-/-^ and wild-type mice. *Vav-*Cre *Ifnlr1^-/-^* mice were generated by crossing *Ifnlr1^fl/fl^* mice with mice expressing Cre recombinase under the *Vav* promoter (Jackson labs 008610) and bred as Cre hemizygotes with Cre maintained on the female breeder.

### Mouse Experiments

Timed pregnancies were set up by exposing females to soiled male cage bedding for 3 days to promote estrus, then housing single pairs of male and female mice overnight (E0), and separating males and females the next morning (E1). Mice were infected by a subcutaneous route in the footpad with 1000 FFU of ZIKV in 50µL. Wild-type, *Ifnlr1*^+/-^, and *Ifnlr1*^-/-^ mice were administered 2 mg of anti-IFNAR1-blocking antibody MAR1-5A3 by intraperitoneal injection (26). For viral load experiments in non-pregnant mice, blood was collected at 2 or 4 days post-infection (dpi) by submandibular bleed, or at 6 dpi by cardiac puncture into serum separator tubes (BD) and serum was separated by centrifugation in a microfuge at 8,000 RPM for 5 minutes. Spleen, brain, and uterus were collected 6 dpi following perfusion with 20 mL of PBS. For weight loss and survival experiments, mice were weighed each day following infection. Pregnant mice were sacrificed at E15 (6 or 8 dpi). Maternal blood was collected by cardiac puncture in serum separator tubes (BD), and serum was separated by centrifugation in a microfuge at 8,000 RPM for 5 minutes. Dams were perfused with 20 mL of PBS then fetal heads, fetal bodies, and their associated placentas, as well as maternal spleen and brain were collected. Fetal tissues were weighed, and total fetal weight was determined by combining fetal head and body weights. Photographs of fetuses and uteruses were taken at time of harvest, and crown rump length was measured using ImageJ (68). For poly(I:C) experiments, 200 µg of low molecular weight poly(I:C) (InvivoGen tlrl-picw) was administered by intraperitoneal injection at the indicated days following mating. At E15, pregnant dams were sacrificed, whole fetuses and their associated placentas were collected and weighed. Implantations with no discernable placentas or fetuses were classified as resorptions.

### RUBV mouse experiments

*Ifnar1^-/-^, Ifnlr1^-/-^, Ifnar1^-/-^ Ifngr1^-/-^* DKO, and wild-type mice were inoculated with 1,000 or 1x10^5^ FFU of RUBV by subcutaneous injection in the footpad or intranasal administration. Weights were monitored for 14 dpi. For viral load experiments, serum and whole blood were harvested 2, 4, and 7 dpi by submandibular bleed into serum separator tubes (BD) and serum was separated by centrifugation in a microfuge at 8,000 RPM for 5 minutes. Mice were sacrificed at 7 dpi, perfused with 20 mL of PBS, then spleens, lung, and brains were harvested.

### Viral Loads

Tissues were homogenized in 600 µL of PBS using a MagNA Lyser (Roche), then 150 µL of homogenate was added to an equal volume of buffer RLT (Qiagen) for RNA extraction. Viral RNA was extracted using a Qiagen RNeasy kit (tissues) or Qiagen viral RNAmini kit (serum). ZIKV RNA was detected by Taqman one-step qRT-PCR using primer probe set: forward- CCGCTGCCCAACACAAG; reverse CCACTAACGTTCTTTTGCAGACAT; probe56- FAM/AGCCTACCT/ZEN/TGACAAGCAATCAGACACTCAA/3lABkFQ on a BioRad (CFX96) using standard cycling conditions. ZIKV genome copies/mL were determined compared to a ZIKV standard curve of 100-fold dilutions of ZIKV-A plasmid (69), or 100 fold dilutions of RNA extracted from viral stock. RUBV viral loads were determined compared to a standard curve made from 100-fold dilutions of RNA isolated from virus stock.

### IFN-λ activity assay

Tissue homogenates and serum were diluted 1:4 in PBS and 20µL was added to 96 well plates. HEK-Blue IFN-λ reporter cells (InvivoGen) were then suspended at a concentration of 2.8x10^5^ cells/mL in DMEM supplemented with 1µg/mL Puromycin, 10µg/mL Blasticidin, and 100µg/mL Zeocin. The HEK-Blue IFN-λ cell suspension was then added to each well of diluted tissue samples and incubated at 37C° for 24h. Then 20µL of the culture media was added to QUANTI-Blue substrate (InvivoGen) for 1.5hr and absorbance was measured at 620nm (bio-tek, epoch). Absorbance readings were converted to concentration using a standard curve of 10-fold serial dilutions of hIFN-λ2 (PBL11820-1) starting at 2500ng/mL, which was run concurrently with tissue samples.

### IFN-β ELISA

Tissues were homogenized in 600µL of PBS using a MagNA Lyser (Roche). Tissue and serum samples were loaded directly onto ELISA plates according to protocol (Biolegend 439407 Legend Max Mouse IFN-B ELISA kit). Absorbance was read at 450nm (bio-tek, epoch).

### Genotyping

*Ifnlr1* and *Sry* (fetal sex) genotypes were determined by PCR on fetal head RNA samples (which contain co-purified genomic DNA), or on DNA extracted from maternal blood and tail samples using the Quantabio supermix and previously described primers: *Ifnlr1* F15- AGGGAAGCCAAGGGGATGGC-3, R15-AGTGCCTGCTGAGGACCAGGA-3, R25- GGCTCTGGACCTACGCGCTG-3) (67), *Sry* F5-TTGTCTAGAGAGCATGGAGGGCCAT-3 and R5-CCACTCCTCTGTGAC ACTTTAGCCCT-3′ (70).

### Viral replication and IFN response assays

Human endometrial stromal cells (HESC-T) were obtained from Dr. David Aronoff (Vanderbilt University). HESC-T were decidualized by culturing cells with 0.5 mM 8-Bromo-cAMP (Sigma B5386), 1 μM medroxyprogesterone acetate (MPA, Sigma M1629), 10 nM 17b-estradiol-acetate (estrogen E2, Sigma E7879) for 5 days as originally described (71). Cells were plated at 500,000 cells/well in 6 well plates and infected at an MOI of 1 with ZIKV (H/PF/2013), DENV4 (TVP-360), or RUBV (M33) in 300µL/well. Supernatant was collected at 4, 24, 48, and 72 hours post-infection, and titered by focus forming assay as described above. A549, JEG3, HTR8, and decidualized HESC-Ts were treated with 50ng/mL IFN-λ (PBL11820-1) or 5ng/mL IFN-β (PBL11420-1), or infected with ZIKV (H/PF/2013) or DENV4 (TVP-360) at an MOI of 1. After 24 hours, RNA was extracted from cell lysates (Qiagen RNeasy kit) and *IFIT1* expression was measured by qRT-PCR (IDT Assay ID Hs.PT.561.20769090.g).

### Statistics

All statistics were performed using GraphPad Prism. Significant differences in fetal weights, viral loads with standard distributions (maternal spleens, placentas), and placental IFN-λ levels were assessed by ANOVA. Significant differences in fetal-head viral loads were calculated by Mann-Whitney.

## ACKNOWLEDGEMENTS

This work was supported by R01 AI139512 (H.M.L.) and start-up funds from UNC Chapel Hill Department of Microbiology & Immunology and the Lineberger Comprehensive Cancer Center. R.L.C. and D.T.P. were supported by T32 AI007419; R.L.C. was supported by a UNC dissertation completion fellowship.

## SUPPLEMENTAL DATA

**Supplemental Figure 1.**
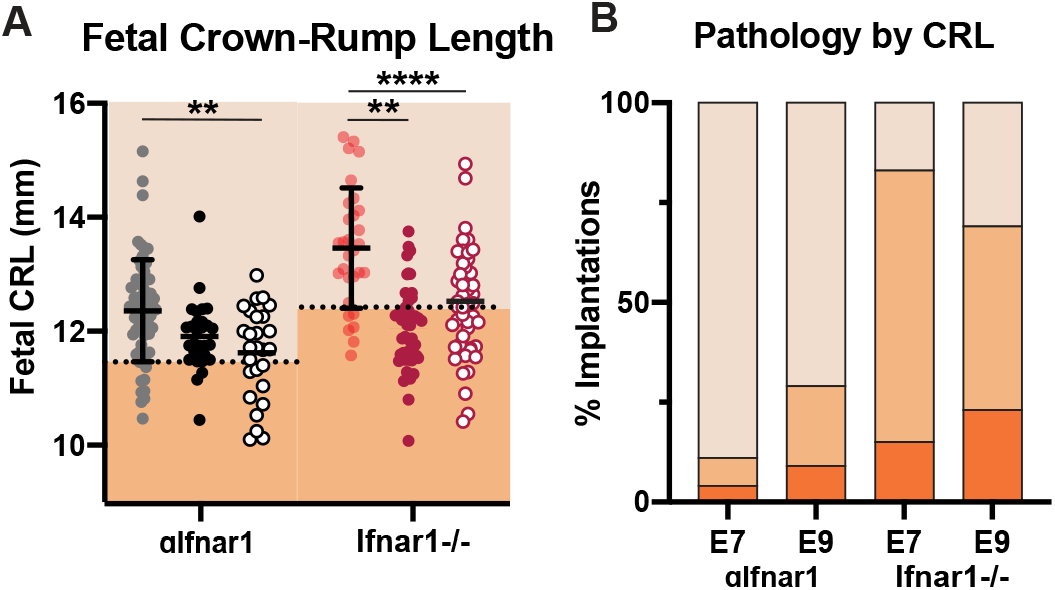
Fetal pathology as assessed by crown-rump length. **A.** Crown-rump length (CRL) of intact fetuses (i.e. not resorbed) from Figure 1 was measured using ImageJ. Fetuses <1 standard deviation from the mean of mock-infected (below dotted line) were classified as having intrauterine growth restriction (IUGR). Intact fetuses with CRLs significantly different from mock pregnancies (calculated by ANOVA) are indicated **** P<0.0001, ** P<0.01 **B**. Proportions of fetuses exhibiting IUGR or resorption.

**Supplemental Figure 2.**
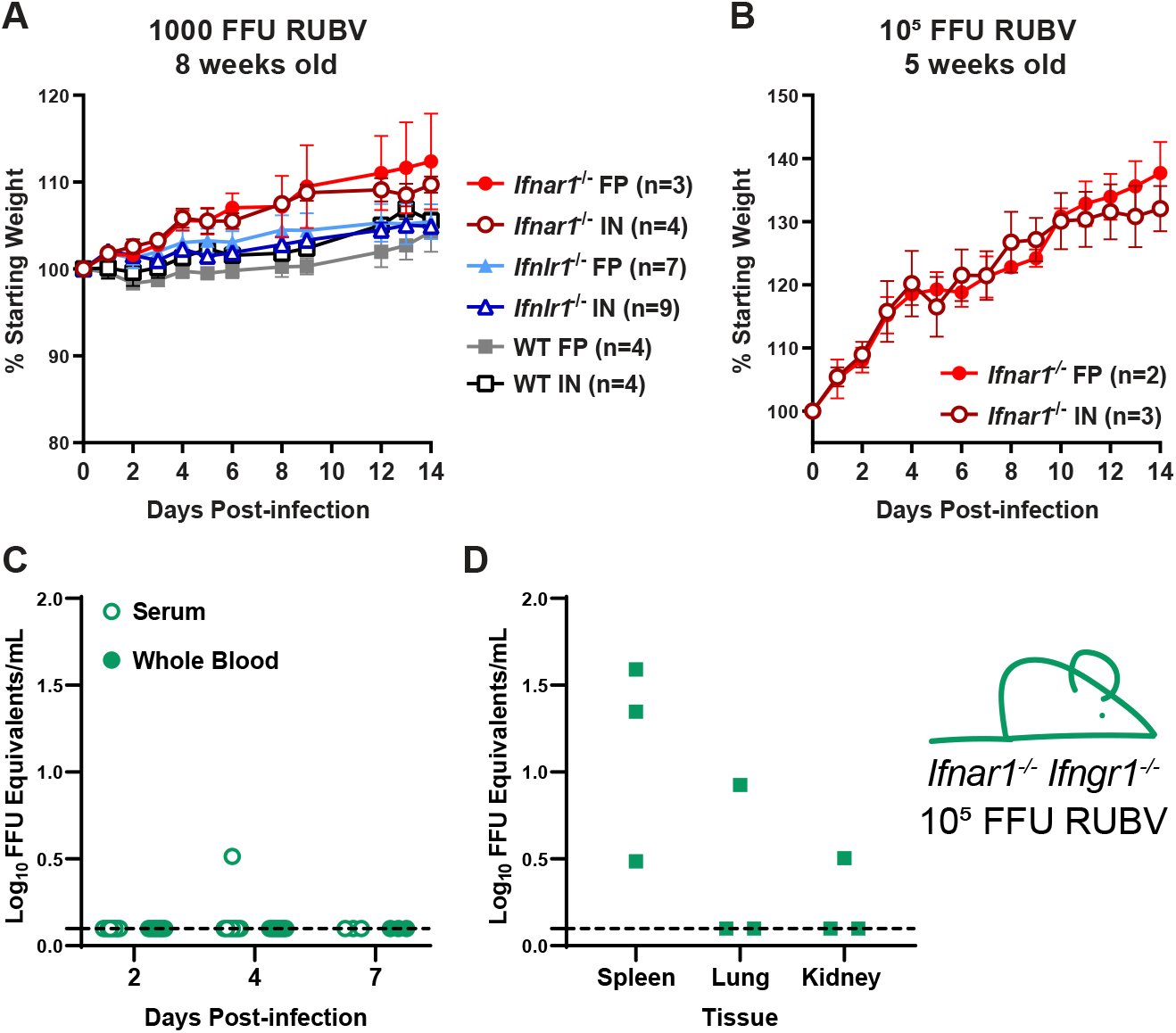
Rubella virus does not cause pathology in mice. **A** and **B.** 8- or 5- week-old male and female WT, *Ifnlr1^-/-^*, or *Ifnar1^-/-^* mice were infected with 1000 FFU or 100,000 FFU of RUBV by subcutaneous inoculation in the footpad (FP) or intranasal inoculation (IN). Weight was monitored for 14 days post-infection and is shown as the mean ± SEM of the indicated number of mice per group. **C** and **D.** 5-week-old *Ifnar1^-/-^ Ifngr1^-/-^* DKO mice were infected intravenously with 100,000 FFU of RUBV. Whole blood and serum were collected 2, 4, and 7 dpi and tissues were harvested 7 dpi. Viral loads were determined by qRT-PCR.

**Supplemental Figure 3.**
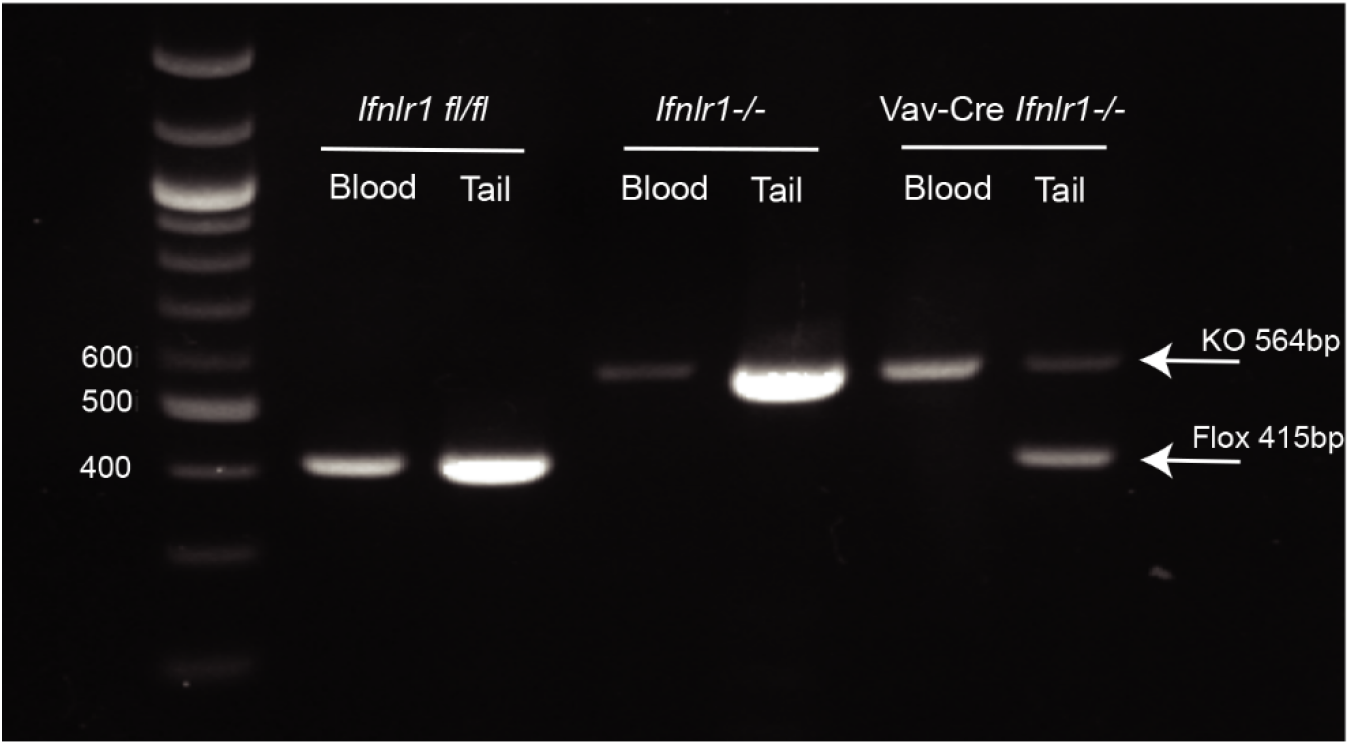
Validation of Vav-cre conditional knockouts. Tails and whole blood were collected from *Ifnlr1^fl/fl^, Ifnlr1^-/-^*, and *Vav*-Cre *Ifnlr1^-/-^* mice and *Ifnlr1* genotype determined by PCR. Knockout band: 564bp, floxed band: 415bp.

**Supplemental Figure 4.**
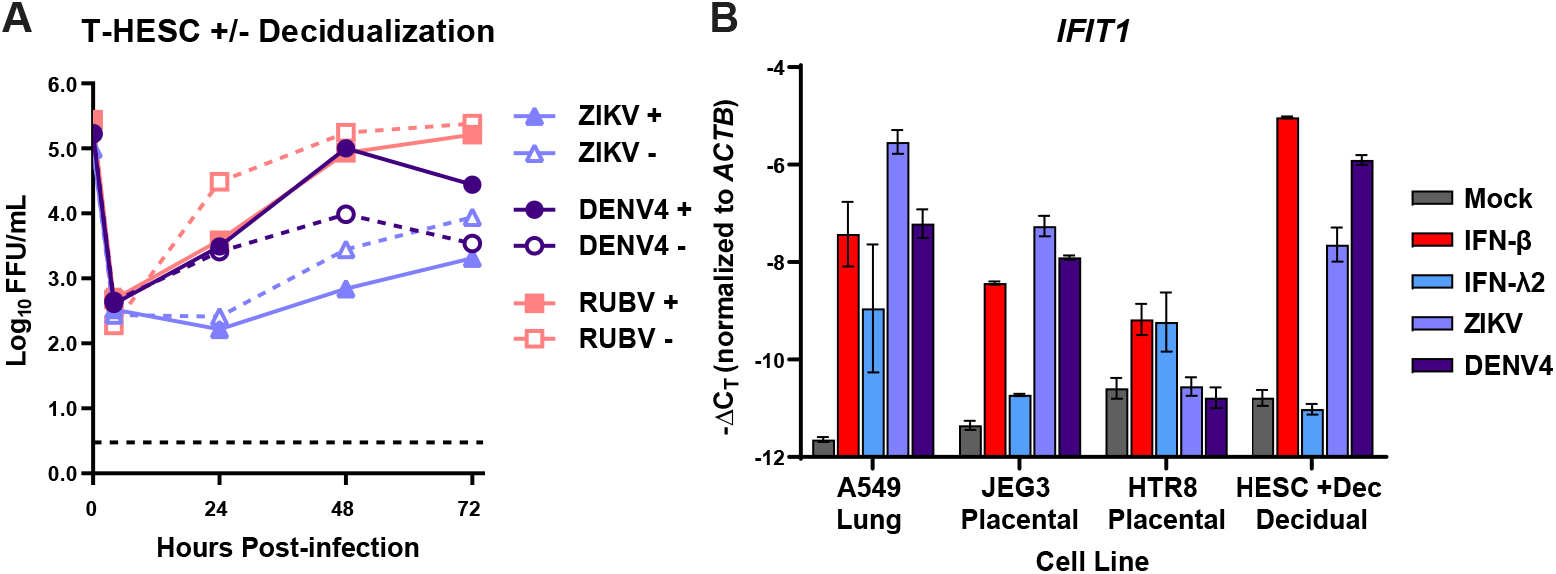
Decidual cell lines are permissive to viral infection, but do not respond to IFN-λ. **A.** Immortalized human endometrial stromal cells (T-HESC) were decidualized (+) or left undifferentiated (-). Decidualized and non-decidualized cells were infected with DENV- 4, ZIKV (strain H/PF/2013), or RUBV at an MOI of 1. Supernatants were harvested at 4, 24, 48, and 72hpi and titered by FFA. **B.** Lung epithelial (A549), placental (JEG3, HTR8), and decidualized human endometrial stromal cells (T-HESC) were treated with IFN-λ (50 ng/mL) or IFN-β (5 ng/mL), or infected with ZIKV (strain H/PF/2013) or RUBV at an MOI of 1. RNA was isolated from cells 24 hours after treatment, and *IFIT1* induction was measured by qRT-PCR.

